# Stepwise assembly and mechanical stability of nsP3 Helical Scaffolds in Chikungunya Virus

**DOI:** 10.64898/2026.11.05.764274

**Authors:** Natalia Martín-González, Andrei Karpov, Juan Reguera, Ignacio Casuso

## Abstract

Chikungunya virus (CHIKV) is a positive-sense, single-stranded RNA virus that replicates its genome through a viral replication complex (RC) essential for successful infection. The RC consists of four non-structural proteins: nsP1, nsP2, nsP3, and nsP4. In this study, we focused on nsP3, which comprises three distinct domains: the alphavirus unique domain (AUD), the long and intrinsically disordered hypervariable domain (HVD), and the enzymatic macrodomain (MD). Notably, nsP3 assembles into helicoidal macrostructures, termed helical scaffolds (HSs), which are components of the CHIKV replication complex and are involved in the formation of macromolecular traps in the cytoplasm of the infected cell called Alphagranules (AGs). HSs form tubular helical scaffolds assembled through the AUD, with the MD and HVD flexibly deployed to the scaffold exterior. Despite their functional importance, the mechanisms governing HS formation remain poorly understood. Using High-Speed Atomic Force Microscopy (HS-AFM) and Electron Microscopy (EM) we achieved new findings on the structure, mechanics and assembly of these macrostructures. Our results show the nsP3 monomers follow intermediate steps before completing the assembly into full helical scaffolds, with each nsP3 domain contributing differently to this process. The macrodomain (MD) provides mechanical stability, yet it fractures under tens of piconewtons of force. This rigidity acts as a mechanical regulator that constrains the position of the HVD, which in turn mediates interactions with host factors and neighboring HSs. Through this interplay, nsP3 achieves both structural cohesion and functional adaptability, promoting the higher-order organization of HSs into honeycomb-like networks.

## INTRODUCTION

Chikungunya virus (CHIKV) is a mosquito-borne pathogen belonging to the Togaviridae family and alphavirus genus. Its infection leads to symptoms such as fever, rash, a strong invalidating arthritis generally lasting for weeks and in severe cases, death ^1^. The first vaccines for CHIKV prevention were recently approved ^2^, however no antivirals are available for the treatment of Chikungunya fever, underscoring the need for a deeper understanding of its replication mechanisms to develop effective therapeutic approaches.

CHIKV undergoes a crucial step during its replication process where double-stranded RNA intermediates emerge. To protect these intermediates from native immune system of the cell, alphaviruses store the dsRNA within membrane spherules gated by a replication pore that allows viral dsRNA synthesis inside and the capped and polyadenylated genomic RNA outside. This pore is composed of four non-structural viral proteins (nsPs) encoded in the first open reading frame of the CHIKV genome. NsP1 acts as the foundation of this pore by forming a membrane-bound dodecameric ring and simultaneously playing a role in capping ^3-4^. Positioned within this ring are nsP4 (viral RNA-dependent RNA polymerase) and nsP2 (viral helicase and protease), both directly involved in viral RNA replication. More mysterious is the role of nsP3; the combination of cryo-electron tomography in infected cells and the recent discovery of nsP3 helical scaffolds (HSs) suggest that nsp3 forms a ring structure in the cytoplasmic side of the nsP1-2-4 complex. This connection highlights the structural and regulatory role of nsP3 in RNA synthesis ^5-7^. The HSs are also found in the cytoplasm of infected cells, forming large cytoplasmic alphagranules by the honeycomb arrangement of multiple parallel HSs. AGs accumulate multiple host and viral factors that in the case of CHIKV include cellular RNA binding proteins as G3BP (essential component of stress granules) or YBX1, membrane and cytoskeleton associated proteins as amphiphising 1 (i.e. BIN 1), CD2BP or FHL1, as well as viral nsP2 protein, capsid protein and genomic RNA^5, 8-11^. Alphagranules thus act as veritable molecular traps which function is important for infection but not fully understood due to their complex composition and the ubiquitous and multifunctional nature of nsP3.

The nsP proteins are produced as a polyprotein which is proteolytically processed into four separate proteins during infection. The crystal structure of the nsP3 was achieved from a Sindbis virus protein construct including the nsP3 alphavirus unique domain (AUD) and N-terminal macrodomain (MD) fused to nsP2 ^12^. In consequence, AlphaFold3 produces very accurate prediction of the CHIKV nsP3 including MD, AUD domains and the N terminus of the HVD (pTM = 0.62, Fig.1.a). The model shows the three domains of nsP3: 1. A N-terminal macrodomain with enzymatic ribosyl hydrolase activity ^13^; 2. An alphavirus unique domain conserved among alphavirus species, and 3. A C-terminal hyper-variable domain which is long and intrinsically disordered and that holds multiple peptide sequences for the specific binding of numerous host factors ^5, 8-11^.

**Figure 1.**
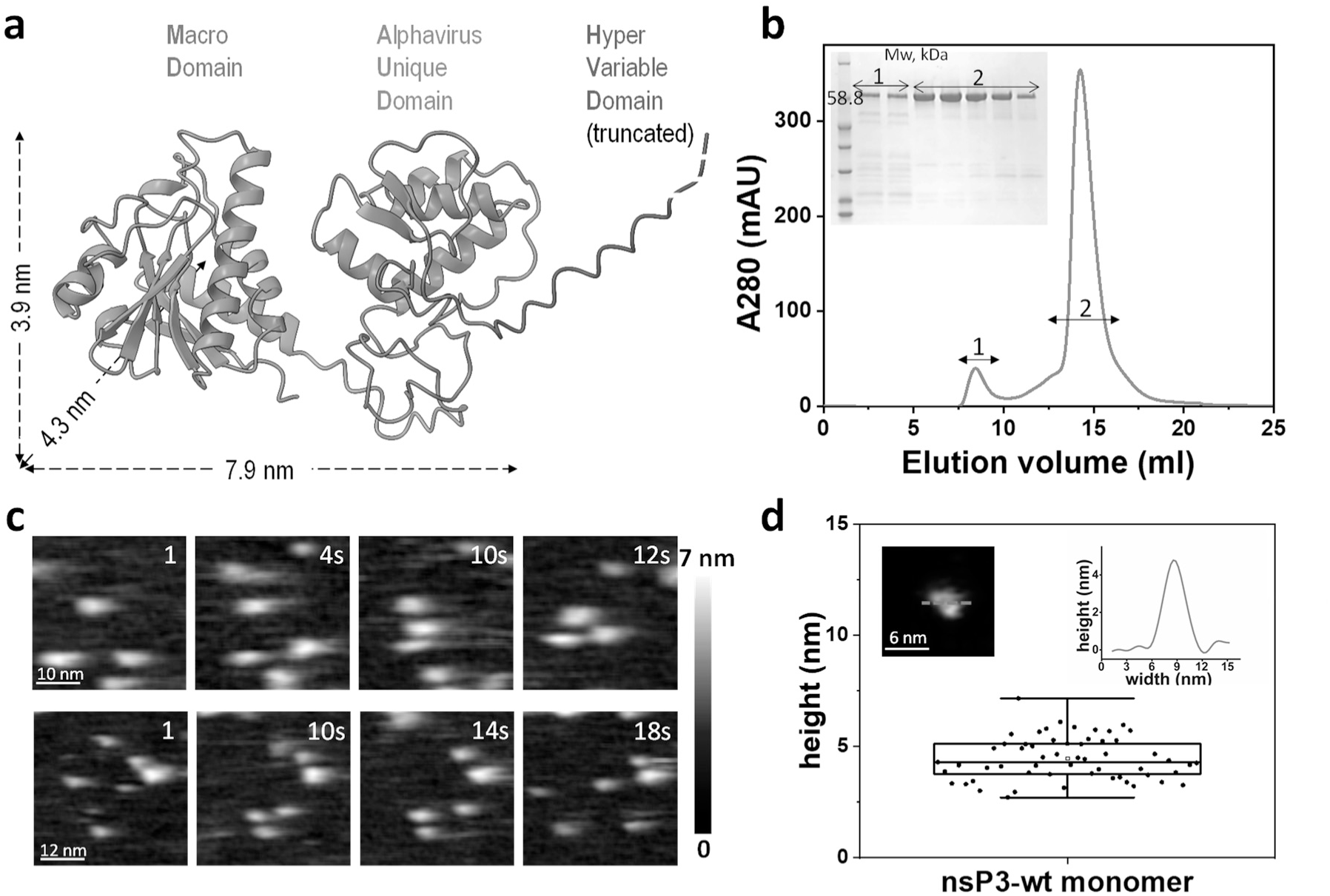
nsP3-wt monomers imaged by HS-AFM are consistent with structural models. **(a)** Structure prediction of nsP3 monomer with truncated HVD by Alphafold3. nsP3 domains are represented in cartoons and shown in different colours: MD in blue, AUD in green and the N terminus of HVD in red. **(b)** Size-exclusion chromatography of nsP3 expressed in insect cells after nickel affinity purification at 500 mM NaCl. Two peaks were observed: (1) high molecular weight species and (2) monomeric nsP3. Inset: SDS-PAGE of fractions from both peaks showing CHIKV nsP3 at the expected 58.8 kDa. **(c)** Representative frames of nsP3 monomers acquired over 60 s under 500 mM NaCl, during which no higher-order assemblies were observed., Slight noise reduction by gaussian blur was applied. **(d)** Height distribution of nsP3 monomers imaged with HS-AFM at 500 mM NaCl. Inset: AFM topography image of a nsP3-wt monomer (top left) and its topographical profile (top right) corresponding to the green line traced in the image.

Most critically, during infection within cells, our previous study (Kril et al. 2024) showed that the nsP3 assembles into tubular structures, lengthening as infection progresses and accumulating in cytoplasmic AGs. Cryo-EM volume reconstructions at 2.3 Å resolution showed that the nsP3’s MD and AUD domains form two-start helical scaffolds. This means that the tubular structure consists of two intertwined helices (termed hA and hB), each following a right-handed path, with approximately 11.4 nsP3 monomers per turn, forming a stable and repetitive filamentous architecture. The inner tube is highly defined by AUD domains which maintain rigid contacts that can be visualised at high resolution by electron microscopy. MD appeared at low resolution at the outer side of the tube showing a highly flexible arrangement of the domains. The HVD domain was not visible due to its intrinsically disordered nature, only few residues emerging from the AUD C terminus to the exterior could be localised at low resolution. The HSs can be assembled and disassembled in vitro by decreasing or increasing NaCl concentrations, respectively ^5^.

The understanding of the nsP3 helical scaffold structure-function relations demands a combination of technical approaches going from biochemical assays to state-of-the-art microscopies; the combination of High-Speed Atomic Force Microscopy (HS-AFM) ^14-15^ and cryo-EM deliver a dataset that uniquely combines dynamic, mechanical and structural information. HS-AFM enables the visualization of flexible and disordered regions that escape cryo-EM averaging, while also probing the mechanical stability of the assemblies. Following the identification of nsP3 helical scaffolds in infected cells by Kril et al. (2024), the present work defines the molecular assembly pathway and mechanical determinants underlying the formation, stability, and higher-order organization of these structures. Here, by integrating dynamic topographical imaging with static structural information, we capture intermediate assembly states, reveal how individual nsP3 domains contribute to filament growth, and probe the mechanical constraints that shape the architecture of the emerging scaffolds. This multimodal approach uncovers structural transitions and force-dependent behaviours that are inaccessible to either technique alone, ultimately offering new insight into the molecular principles governing HS formation and higher-order organization.

## RESULTS

### Stable Monomeric nsP3 Conformation Observed by HS-AFM is Consistent with Predicted Structural Dimensions

The nsP3 protein was expressed in insect cells and purified by serial affinity and size exclusion chromatography at 500mM NaCl. The size exclusion chromatography showed two peaks: The first peak corresponds to nsP3 assemblies, and the second peak is consistent with the monomeric nsP3 (58.8 kDa) (Fig.1.b). For HS-AFM exploration, the monomeric nsP3 was absorbed on the mica surface at the same salt concentration used during purification. The HS-AFM first showed the nsP3 individual globular proteins diffusing on the mica surface (Fig.1.c), hence indicating low surface-protein interaction (Suppl. Movie 1, Suppl Fig. 1). During the 407 s observation period, no higher-order assemblies were detected under these high-salt conditions. Topographical characterization of the monomers, obtained by tracing topographical profiles (height measurement accurately represents the protein diameter, whereas AFM-measured lateral width is prone to dilation of the observed protein diameter^16^) revealed a height distribution of 4.5 ± 1.0 nm (mean ± SD, N = 57) (Fig.1.d). This measured height closely matched the molecular model predicted by AlphaFold3 (Fig.1.a) in a configuration where the protein lies flat on the surface.

To complement the surface-based HS-AFM observations with bulk measurements, we performed dynamic light scattering (DLS) experiments on nsP3-wt under high- and low-salt conditions (Supp. Fig. 2). Despite the limitations of DLS, including the low signal-to-noise ratio for small proteins and the influence of the helical rod-like structure of nsP3 HSs (which deviates from the spherical particles that are optimal for DLS measurements), we were consistently able to detect that at 500 mM NaCl, nsP3-wt displayed a stable hydrodynamic radius (Rh) of 6.0 ± 0.5 nm over the entire measurement period (∼1 h), consistent with a monomeric state. This value agreed with the monomer height measured by HS-AFM. In contrast, upon dilution to 50 mM NaCl, the first measurement (∼2.5 min after dilution) already revealed a large Rh of 40 ± 3 nm, which remained constant thereafter. These results indicate that nsP3 rapidly assembles into large supramolecular structures within minutes after salt reduction, consistent with the fast helical scaffold formation observed by electron microscopy in (Kril et al., 2024) ^5^.

**Figure 2.**
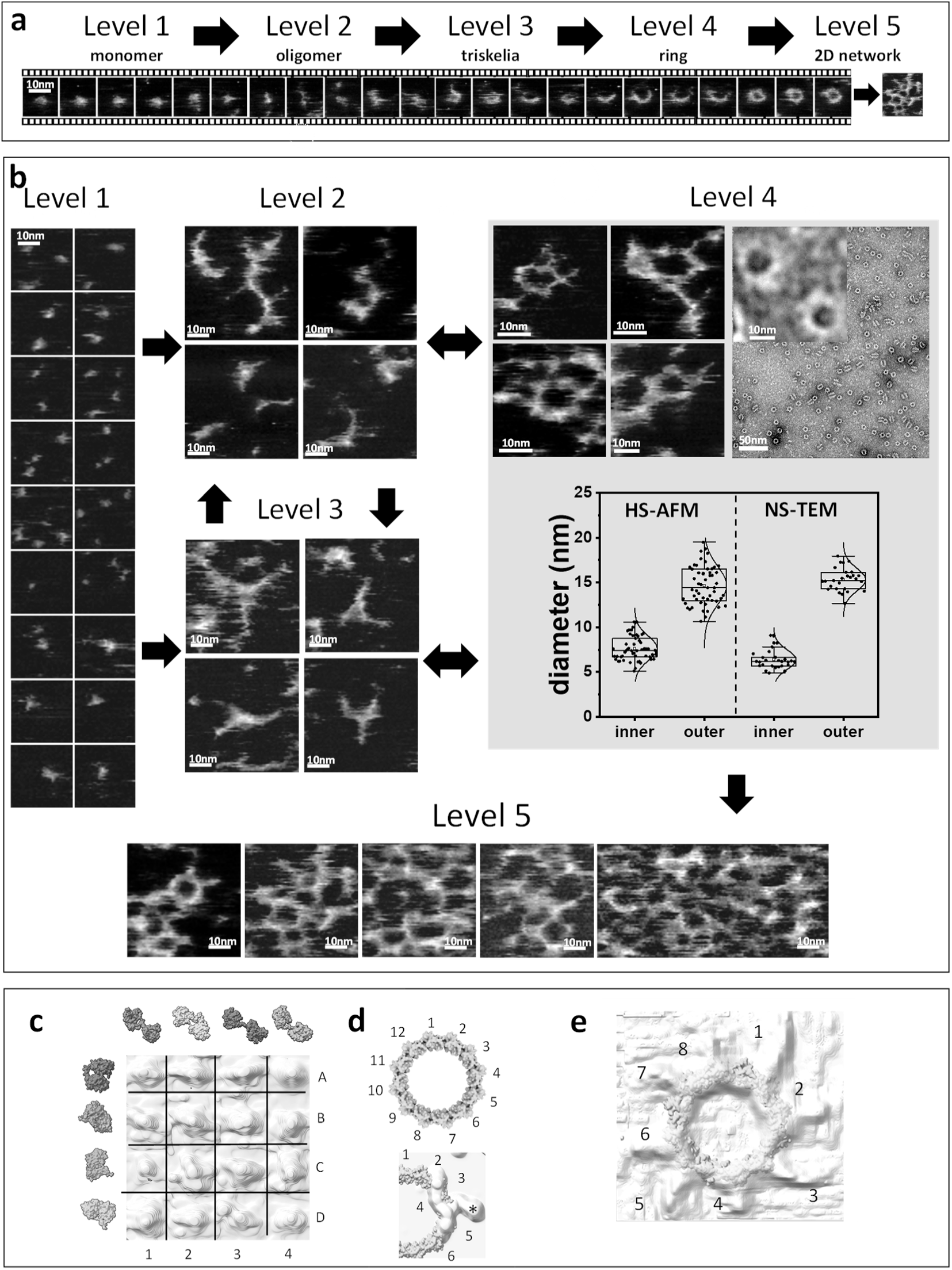
Incremental hierarchical organization of nsP3 during early stages of Helical Scaffold formation visualized by HS-AFM and NS-TEM at 50 mM NaCl. **(a)** Definition of the five different levels or organization of nsP3. From left to right, i) monomer (isolated nsP3 molecule), ii) oligomer (small associated assembly without a defined higher-order geometry), iii) triskelion (threefold branched assembly), iv) ring (closed circular assembly with a central channel) and v) 2D network (extended interconnected assembly). Representative HS-AFM time series showing the dynamic progression from monomer to ring through the different levels of nsP3 organization. Consecutive frames are separated by 1 s. **(b)** HS-AFM images of the five levels of nsP3 monomers assembly. Bidirectional arrows indicate reversible transitions between different levels observed in the HS-AFM movies. Level 4, shows a comparison of HS-AFM image (left) with a NS-TEM image that shows ring like structures (right). Comparison of inner and outer ring diameters measured by HS-AFM and NS-TEM showed no significant differences between the two populations. **(c)** Egg box representation of AFM volumes corresponding to nsP3 monomers showing conformational heterogeneity but volume size consistency. On the left in green are represented the surfaces of four alphafold predictions of nsP3 at the same size-scale than the AFM volumes. On the top and with different colors are shown four models generated by AFlex from alphafold model in dark green. AFlex models are extended showing clearly the two domains of nsP3 (MD and AUD) wilst the alphafold models are more compact. The volumes also show two lobes (3D, 2D, 4C) or single lobes extended (2A, EA) or more compact (4A, 4B). **(d) Top**, circularized model of 12 nsP3 AUD domain rings, the MD and HVD are absent in the model, the 12 subunits are shown. **Bottom**, superposition of a triskelion with the nsP3 12-fold ring showing four subunits (2-5) covered and an extra density labeled with an * which could correspond to macrodomains radially projected. **(e)** Superposition of an AFM-topography ring and the 12-fold AUD ring. Eight radial densities appear with different sizes, they are numbered (1 to 8). Two small radial densities (4, 5, 7 and 8) and four larger (1, 3, 6, 2) are detected, these could correspond to MD and HVD domains, which may establish flexible intra-ring (“cis”) and inter-ring (“trans”) interactions on the ring surface on the ring surface, consistent with the interactions revealed by the cryoEM structure of the nsP3 helical scaffolds reported in Kril et al. 2024 ^5^. Here, the terms "cis" and "trans" refer to interactions occurring within the same ring and between neighbouring rings, respectively.. The planar restrain induced by the mica would prevent these interactions from growing in the Z axis being thus projected radially whether as interacting MD dimers or monomers (e.g. 4 dimers and 4 monomers = 12 monomers).

### Hierarchical Assembly of nsP3 Monomers into Helical Scaffolds Revealed by HS-AFM

Reducing the salinity to 50mM NaCl, HS-AFM movies revealed dynamic protein-protein interactions among nsP3 monomers following the initial globular protein shape (Supp. Movie 1 2 and 3). A taxonomy of structures of the nsP3 is depicted (Fig. 2.a), among them, the nsp3 rings that are observed by negative-stain transmission electron microscopy (NS-TEM (Fig. 2.b, level 4 right), indicating that both techniques provide compatible readouts of the structures assembled by nsP3. However, no earlier assembly stages than the ring could be identified by NS-TEM. By HS-AFM we analyzed approximately a quarter of a million interacting nsP3 assemblies across 7,500 frames and we could identify five incremental levels of hierarchical organization not observed by NS-TEM (Fig. 2.a). The first level, the monomers (Fig.2.a, Fig.2.b, level1) adopted higher conformational heterogeneity than that at 500 mM NaCl, including single and double lobes structures (Fig. 1.c-d), but volume size consistency (Fig.2.c, comparison of an egg box representation of different shapes derived from HS-AFM observations with similar AlphaFold and AFlex models obtained in silico). Besides the monomers, the HS-AFM found other intermediate levels of organization: the oligomers assembled to form two dominant types of architectures: the arches (level 2) and the triskelia (level 3) (Fig.2.a, Fig.2.b, Supp. Fig 3.a, and Supp. Fig 3.b). The HS-AFM movies show that these intermediate structures constantly interconvert, dynamic exchanging shapes between them. The hierarchical levels differ in their lifetime, the nsP3 is mostly found in the form of intermediate and higher levels. Most of the nsP3 is found assembled in intermediate hierarchical levels, only a fraction (>1% in volume) is found in the form of isolated monomers (Suppl. Movie 2), hence, indicating that the monomers fuse with high affinity, being the triskelia and rings the building blocks more significant in the construction of the helical scaffolds. We found that the 180 degrees threefold geometry of the trikelia is a precursor to higher levels of organization in the nsP3 assembly. When the local density of nsP3 proteins reached sufficient levels, a higher organization emerged often by the interaction of triskelia; closed ring-like structures appeared (Fig. 2.b, level 4). Superposition of the topography of the triskelia with four subunits of a circularized model of nsP3 AUD domain rings shows that an extra density which could correspond to triskelion’s macrodomains radially projected when they form the ring structure (Fig.2a, Fig. 2d bottom). The closed structures arise from the interaction of preformed arches and triskelia (Suppl. Fig. 3b), whereas direct formation of ring-like structures from isolated monomers was not observed. Importantly, the diameters of the closed structures correlate well with the nsP3 rings that were visualized by NS-TEM;Fig.2a, Fig. 2b, level 4). For HS-AFM, the inner and outer diameters were measured from four radial topographic profiles at equivalent height levels, as illustrated in Suppl. Fig. 4. The inner and outer diameters were 8 ± 1 nm and 15 ± 2 nm, respectively, by HS-AFM (mean ± SD, N = 54), and 7 ± 1 nm and 15 ± 1 nm, respectively, by NS-TEM (N = 27).The values did not differ significantly between techniques (one-way ANOVA, p > 0.05), suggesting that arches and triskelia, although not visible in NS-TEM images, are likely present in the solution imaged by NS-TEM and that these early interactions are decisive for the subsequent events leading to the formation of helical scaffolds.

**Figure 3.**
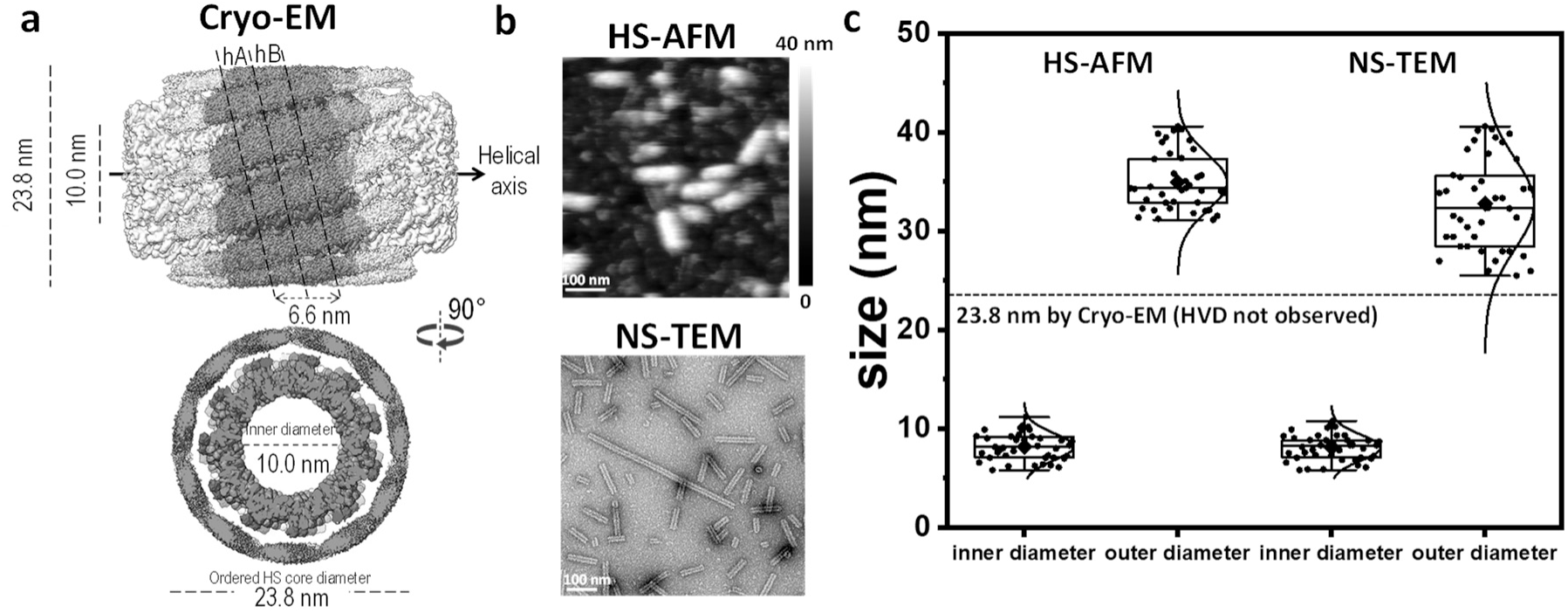
Recombinant nsP3-wt. Last stages of assembly into helical scaffolds. **(a)** Cryo-EM structure of nsP3 helical scaffold assembled in vitro from purified monomers in axial and lateral projections. Note that there is an interstitial space between the AUD colored in green and the MD colored in blue which could be filled by the disordered HVD which is visible only at its N terminus (colored in red). **(b)** HS-AFM (top)and NS-TEM (bottom) representative image of HSs population. **(c)** Helical scaffold inner and outer diameter analysis from HS-AFM and NS-TEM images. Inner and outer diameters were quantified from cross-sectional profiles using technique-specific criteria. In HS-AFM, the inner diameter was measured as the lateral distance between equal height points delimiting the exposed channel in partially disassembled HSs, while the outer diameter was obtained from intact HSs using equivalent height-based edges (cartoon at Fig. 4c). In NS-TEM, diameters were extracted from intensity profiles, with the inner diameter defined by uranyl acetate penetration into the channel and the outer diameter by the full contrast-defined filament width. The 23.8 nm diameter measured by cryo-EM reflects the ordered HS core, as the HVD domains are not resolved due to their intrinsically disordered and highly dynamic nature.

The HS-AFM movies show further levels of complexity unseen by NS-TEM: at the fourth level, successive incorporation of nsP3 monomers into polygons led to the emergence of rectilinear branches from the polygon angles (Fig. 2.a level 4, and Fig. 2.e). By overlaying an AFM-topography ring with the 12-fold AUD ring, as it could be assembled in replication complexes (Kril et al. 2024 ^5^), eight distinct radial densities of varying sizes were identified and numbered (1 to 8, Fig. 2.e). Among these, four smaller densities (4, 5, 7, and 8) and four larger ones (1, 2, 3, and 6) were detected. These densities may correspond to MD and HVD domains, which appear to interact flexibly both within the ring (in cis) and between rings (in trans) on the ring surface. Such interactions mirror those described in the cryo-EM structure of nsP3 helical scaffolds, as reported by Kril et al. 2024 ^5^. In addition to the circular geometry, the subsecond imaging of the HS-AFM observed temporary transitions to other non-circular closed structures, such as regular pentamers and hexamers, which dynamically fluctuated between them (Supp. Fig. 3.c), as the arcs and triskelia did, confirming overall structural variability; this interconversion is presumably driven by the incorporation and removal of nsP3 monomers.

A fifth level emerges when branched polygons interacted assembling a two-dimensional (2D) network that progressively expands on the surface (Fig. 2.b level 5). The 2D network was unstable and underwent continuous dynamic reassembly (Suppl. Fig. 5.a). Presumably, the interaction of nsP3 with the underlying mica surface favors the assembly into a flat geometry, that hinders the formation of HSs, never observed under these conditions. On the mica, the latest stage of the HS formation that assembles is that of the 2D network (level 5). The observation of the 2D network demonstrates that intermediate hierarchical levels exist between the nanometric lower levels (triskelia and rings) and the final HS micron-construction. In solution, these intermediate honeycomb-like assemblies constituted of interacting rings and triskelia (Level 5) would bend into cylindrical geometries.

In the HS-AFM movies, all five stages of nsP3 incremental assembly shared the same height (∼4.5 nm) (Suppl. Fig. 5b). It indicates that these assemblies consist of a single protein layer, finding consistent with cryo-EM data which identifies the HS structures as single-layered.

We conclude that nsP3 building blocks hierarchically assemble into intermediate structures, which in turn assemble to form HSs, and that the assessment of the full pathway of assembly from nsP3 monomer to HS was only possible through the complementary use of HS-AFM and cryo-EM.

### HS-AFM Reveals the Flexible HVD Envelope of the nsp3 Helical Scaffold

Our previously published structure was obtained from HSs purified from insect cells ^5^. Here, using purified nsP3, we determined the structure of in vitro-reassembled helical scaffolds by cryo-EM (Fig 3.a). The structure of the reconstituted nsP3 HSs was resolved at 2.7 Å resolution (FSC 0.143, EMD-56017, Table 1). It showed low-resolution density for the MD, similar to that of the HSs purified from cells, so we consider the purified protein representative of the assembling in cells. As in the case of the HSs purified from insect cells, the HVD region was not observable by cryo-EM due to its intrinsically disordered structure, which fluctuates over time and is averaged out during image processing: In the HSs of purified protein, the diameter of the HS (23.8 nm) does not include the HVDs and possibly part of the MD.

**Table 1.** Cryo-EM data collection, processing, and model refinement statistics.

| Sample | **nsP3-wt** | **nsP3-ΔMD** |
|---|---|---|
|  | PDB: 9TJR | PDB: 9TJS |
|  | EMD-56017 | EMD-56018 |
| Grids | QF R2/1 300 mesh (Au) | QF R2/2 300 mesh (Au) |
| Vitrification method | Leica GP | Vitrobot |
| Microscope | Glacios 200kV | Glacios 200kV |
| Magnification factor | 160000 | 130000 |
| Detector | Falcon 4i | Falcon 4i |
| Recording mode | Counting | Counting |
| Electron dose rate (e<sup>-</sup>Å<sup>-2</sup>s<sup>-1</sup>) | 17.4 | 17.4 |
| Total electron dose (e<sup>-</sup>Å<sup>-2</sup>) | 40 | 40 |
| Pixel size (Å) | 0.71 | 0.90 |
| Number of frames | 40 | 40 |
| Total exposure time (s) | 2.3 | 2.3 |
| Set defocus range (µm) | from -0.6 to -3 | from -0.6 to -3 |
| Number of micrographs | 7168 | 4610 |
| Number of picked particles | 1073295 | 788186 |
| Number of particles used for refinement | 338857 | 325402 |
| Symmetry imposed | Helical, C1, twist: -164.170°, rise (Å): 2.900, symmetry order: 10 | Helical, C1, twist: -164.170°, rise (Å): 2.891, symmetry order: 8 |
| Resolution (Å) (half-map half-map, FSC threshold 0.143) | 2.7 | 3.3 |
| RMSD bonds (Å) | 0.004 | 0.004 |
| RMDS angles (°) | 0.748 | 0.875 |
| Ramachandran favoured (%) | 100.00 | 98.21 |
| Ramachandran outliers (%) | 0 | 0 |
| Rotamer outliers (%) | 1.49 | 1.49 |
| Map-model cross correlation (CC, main chain) | 0.89 | 0.89 |
| Clashscore | 3.57 | 6.30 |
| MolProbity score | 1.28 | 1.48 |
| Map-model FSC (Å) (threshold= 0.5) | 3.0 | 3.5 |

To complement the cryo-EM data with HS-AFM images of the helical scaffolds, we circumvented the two-dimensional constraints imposed by mica-surface adsorption hindering HS formation, through a two-step process: First, HSs were assembled in solution, next, they were adsorbed onto a flat mica substrate (Fig. 3.b, top). This approach preserved the native HS structure, as confirmed by comparison of the HS-AFM images with solution-phase HSs visualized via NS-TEM (Fig. 3.b, bottom). The lengths of the HSs remain consistent across techniques and preparations (Suppl. Fig. 6).

The HS-AFM topography of the HSs did not exhibit the helicoidal shape seen in cryo-EM. The topography of the HS surface appeared as a furry envelope absent of well-defined structural motifs. This type of AFM image is characteristic of intrinsically disordered proteins ^17^. This suggests that, during imaging, the HS-AFM tip contacted and contoured the soft and flexible outer layer formed by the HVD chains surrounding the MD-AUD core of the HS.

The presence of a disordered "furry" HVD envelope surrounding the helical scaffold (HS) was confirmed through comparative HS-AFM-NS-TEM diameter measurements. For intact HSs, we determined the outer diameter using equivalent height-based edges. HS-AFM revealed an outer HS height of 34.7 ± 2.7 nm (mean ± SD, N = 41; Fig. 3.c) on mica, which is approximately 10 nm larger than the 23.8 nm diameter resolved by cryo-EM (N = 363,876). Using HS-AFM, we measured the inner diameter as the lateral distance between equal-height points of exposed channels that became accessible following partial scaffold disruption (see disassembly assays below).The inner diameter obtained was 8.1 ± 1.4 nm (mean ± SD, N = 41). Moreover, this observation was further supported by NS-TEM, which identified inner and outer diameters of 8.1 ± 1.3 nm (N = 41) and 32.8 ± 4.8 nm (N = 41) respectively (Fig. 3.c). The outer diameter, visualized through staining contrast, reflects the presence of an extended peripheral HVD layer, while the inner diameter remains invariant across techniques due to the structural stability of the AUD core, consistently measured at 8 nm by NS-TEM and HS-AFM.

These results demonstrate that the expanded diameter detected by HS-AFM arises from the disordered and highly dynamic HVD chains, which protrude as a distinct "furry" envelope surrounding the HS core. This envelope not only defines the spatial extent of the scaffold but also likely governs its interactions with host factors, highlighting its functional role in cellular processes.

### Disassembly Assay Reveals a Sequential MD–AUD Core Disassembly following HVD Removal

The force applied by HS-AFM tip causes progressive disassembly of the helical scaffold. We subsequently investigated the mechanical stability of the HSs from HS-AFM movies (Supp. Movie 4). Similar AFM-based mechanical disassembly approaches have previously been used to investigate the mechanical properties of intact viral particles ^18-20^. Here, we apply this approach for the first time to nsP3 HSs. The HS-AFM movies are consistent with transient, non-specific interactions between the AFM tip and protruding HVD domains at the HS surface, resulting in their progressive displacement or removal. With no observable spatial order, the movies show that the initial pulling and removal of an HVD occurred at arbitrary positions along the HS. No noticeable preference of the position where the peeling of the HS starts was found (Fig. 4.a right and Fig. 4.b, the tip pulls out the HVDs at random locations signalled by arrows).

**Figure 4.**
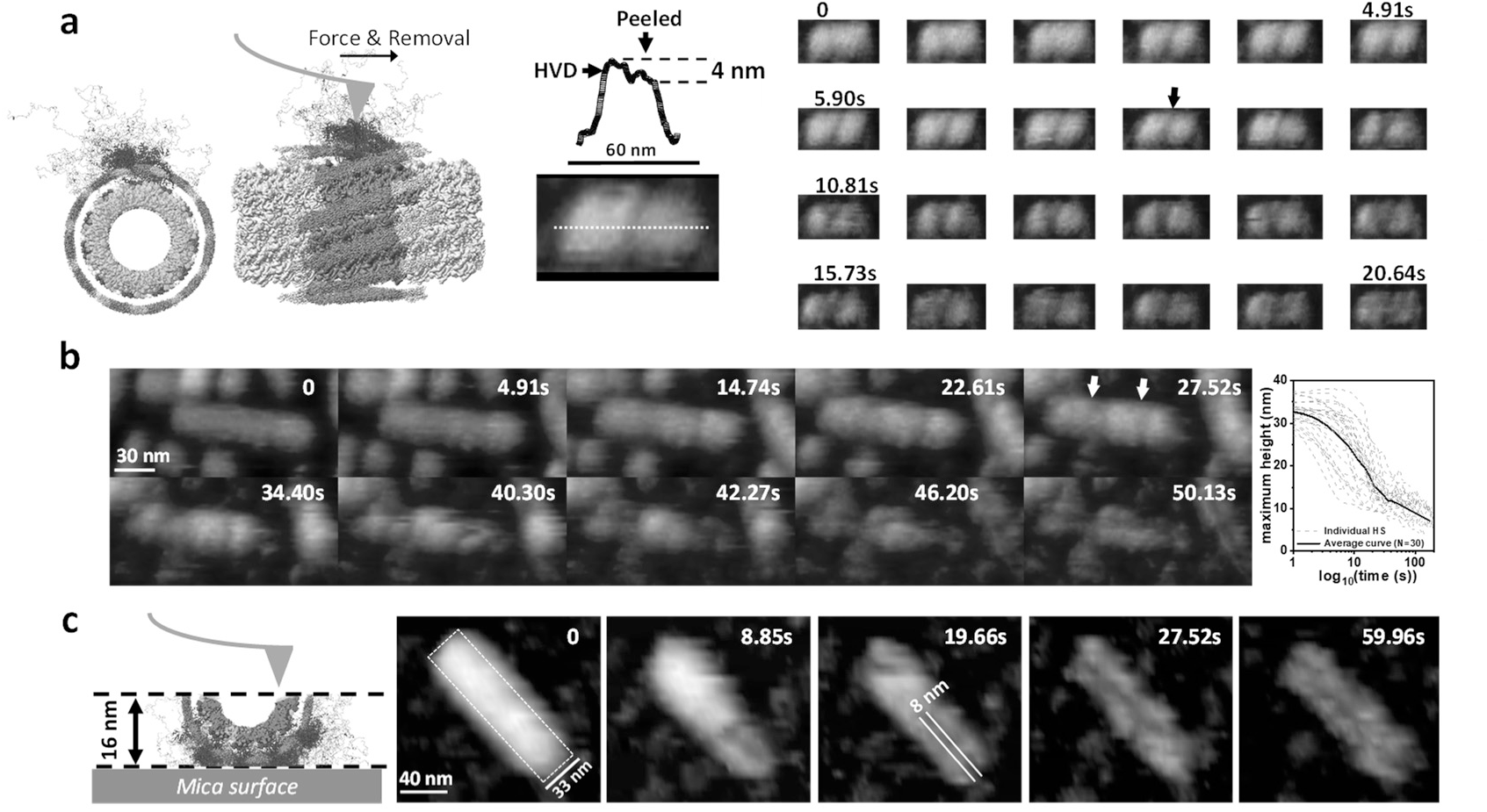
Mechanical disruption of Helical Scaffolds. **(a)** Left, scheme of the AFM tip interacting with HVD domain of the HS. Centre, profile of a HS where the HVD layer has been partially removed on the right side, the topographic profile shows a height step of 4 nm corresponding to the HVD layer. Right, sequence of images showing the progressive removal of the HVD layer by the HS-AFM tip. **(b)** Left, second example of disassembly of the HVDs when pulled by the HS-AFM tip. The white arrows show that the initial pulling and removal of an HVD occurred at arbitrary positions along the HS, which creates a weak point in the HS structure from which disassembly propagates to the neighboring domains. Next, the core of the HS disassembles under the taps of the tip, the structure of the MD and AUD domains fall apart. The lower section of the HS remain as an elongated U-shaped strip, the cumulative height of the AUD, MD, and HVD is 16 ± 2 nm (N=30). **Right**, height evolution along time (in log scale) in N=30 HSs, average curve (black) and propagation speed estimated by the linear fitting of the average black curve, shows that the cooperative disassembly of helical scaffold progresses at a speed of 0.7 ± 0.2 nm/s. **(c)** Left, cartoon showing an HS-AFM tip on a peeled HS how the inner diameter of the HS is measured in the remaining on the mica of the disassembled HS, the full data set is plotted in Fig. 3b. Right, example of inner diameter assessment in a sequence of images of the disassembly.

The movies showed that once the first HVD was removed at a given location, this site became a weak point in the HS structure, from which disassembly propagated through the progressive removal of neighboring domains. (Fig. 4.a right, the removal of the HVDs by the AFM tip expands from the initial weak point). Importantly, the diffuse HVD envelope progressively disappeared during HS disassembly, following the exposure of the ordered MD–AUD helical core. This transition showed regular features across the HS disassemblies (Suppl. Fig. 7): Regions retaining the HVD envelope showed a topographic height approximately 4-5 nm greater than regions where the envelope had been removed (Fig. 4a, centre; Suppl. Fig. 7), consistent with our previous estimates of the HVD envelope. This reproducible transition supports the absence of imaging-related distortions. Eventually, when the HVD envelope was completely removed from sections of the HS, the AFM tip contacted and revealed the characteristic helical outlines imaged by cryo-EM (Fig. 4.a and 4.b).The apparent lateral width of the HS may be broadened by tip-convolution effects and a certain degree of parachuting, particularly in lower topographic regions (Fig. 4c).

Once the HVDs were removed, the core MD-AUD structure of the HS also disassembled: the structure of the MD and AUD domains fell apart progressively, the HS could not maintain its structural integrity, crumbling under the taps of the HS-AFM tip (Fig. 4.c, sequence of HS disassembly). The minimal force required for HS-AFM observation (tens of pN, ^21^) was sufficient to disassemble the HS. The full disassembly process lasted between 10 and 30 seconds (at imaging rates of 1 frame/second), with an axial propagation speed of 0.7 ± 0.2 nm/s (Fig. 4b right, derived from the linear slope of the height–time curve). After disassembly, the lower section of the HS remained as an elongated U-shaped strip on the mica, the three domains were stacked on top of each other. The cumulative height of the AUD, MD, and HVD in this strip was 16 ± 2 nm (N=30), which is thicker than the 13 nm measured by cryo-EM that excludes the HVD. The result provides additional evidence that the HVD layer contributes approximately 4–5 nm to the scaffold thickness, consistent with the independent estimates obtained from the HS-AFM/cryo-EM diameter comparison (Fig. 4.c). Moreover, the U-shaped lower section of the HS enabled the measurement of the inner diameter by HS-AFM (Fig. 3.c). As a lateral measurement, the inner and outer diameter are subject to tip-convolution and it may be slightly overestimated, however, our measurements at the same height and under identical imaging conditions allow us reliable comparison between HSs. These measured values compare well with those obtained by NS-TEM (Fig. 3.c), further supporting the compatibility of both approaches.

Overall, our results confirm the general architecture of HSs, accounting for both ordered and disordered regions. The disassembly assay reveals a progressive disassembly pathway in which HVD removal precedes the disassembly of the underlying MD–AUD core. These findings highlight that the MD and HVD may play roles beyond their respective functions in ADP de-ribosylation and host factor interactions.

### Nsp3 MD, HVD, and AUD Domains Play Distinct Roles in the Hierarchical Assembly of nsP3

To investigate in detail the functional contributions of the MD and HVD domains to the assembly process of the HS, we generated three truncated variants: nsP3 lacking the HVD (nsP3-ΔHVD), nsP3 lacking the MD (nsP3-ΔMD), and a construct consisting solely of the AUD (nsP3-AUD).

While nsP3-wt and nsP3-ΔMD were purified with comparable efficiency, the yield for nsP3-AUD was significantly lower. Notably, nsP3-ΔHVD precipitated immediately following Ni-affinity chromatography, which reduced the concentration of protein available for the size exclusion purification step and in consequence, the final pure protein concentration for the EM and AFM studies. These findings suggest that the HVD is essential for maintaining protein stability in solution.

At 500mM NaCl, at the initial phases of HS assembly on mica, we found that two of the constructs, nsP3-ΔHVD and nsP3-ΔMD, exhibited globular monomeric forms with diffusive movement. Within the limits of our detection, these mutants showed no discernible differences from the nsP3-wt at the globular protein stage (Fig. 5.a). In contrast, the third construct, comprising only the AUD, shows a high tendency to aggregate, making it hard to distinguish individual monomers (Fig. 5.b). These findings indicate that the presence of either the HVD or MD domains limits the aggregation of nsP3, highlighting the critical role of these partner domains in the assembly process from its earliest stages.

**Figure 5.**
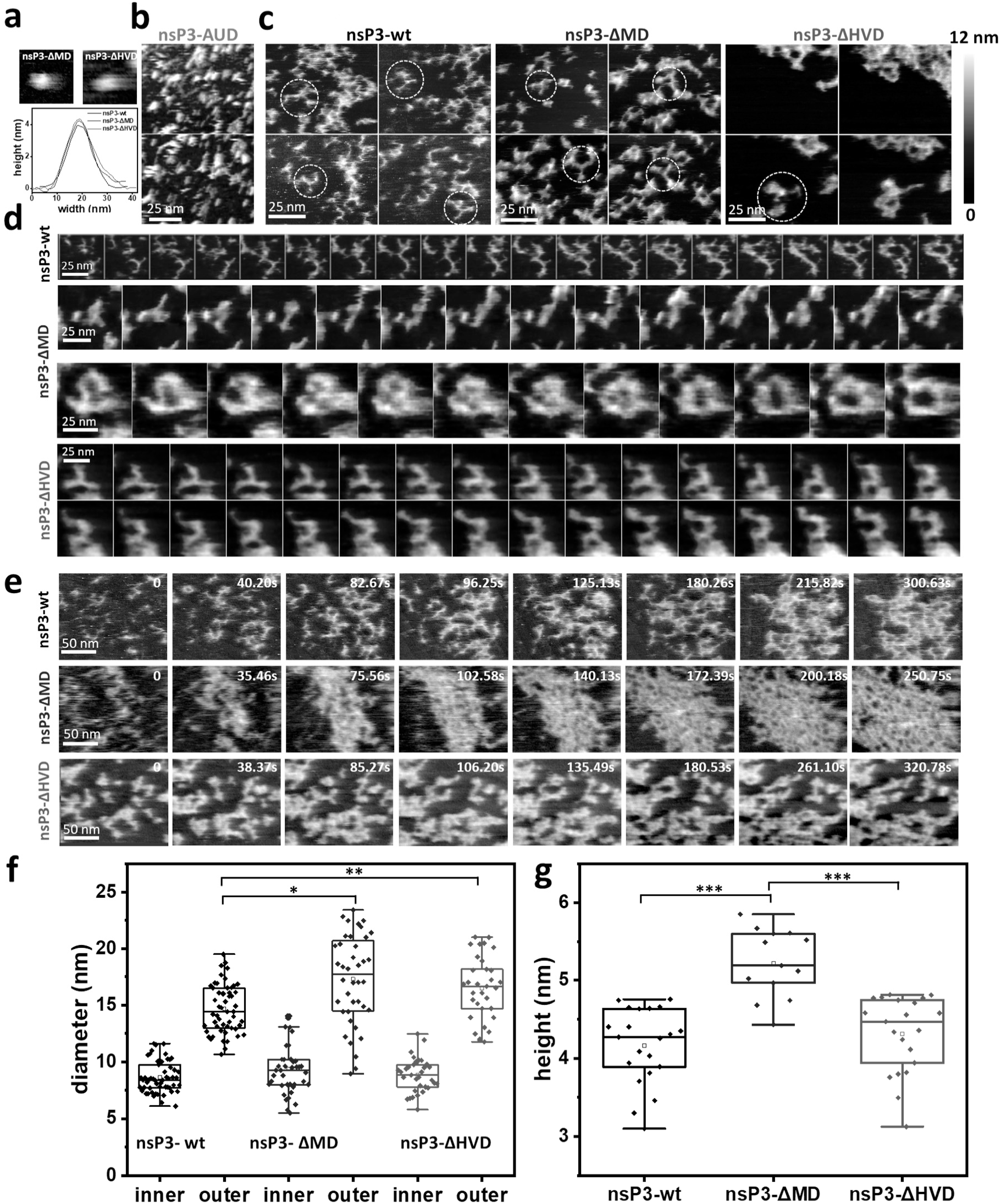
Recombinant constructs of nsP3. First stages of the assembly. Contribution of each domain. **(a)** Top, globular nsP3 monomers of nsP3-ΔMD (left) and nsP3-ΔHVD (right). Bottom, topographical profiles. **(b)** nsP3-AUD images showing that, in this case, the assembly is amorphous and heterogeneous. **(c)** Triskelia of the three (black, blue and red) constructs studied. The nsP3-wt, nsP3-ΔMD and nsP3-ΔHVD images show the population of triskelia, the white dash circles identify the triskelia. From left to right, a decreasing population of triskelia in nsP3-ΔMD and nsP3-ΔHVD mutants is observed (see Suppl Figure 8). **(d)** Dynamics of triskelia assembly into rings. nsP3-wt shows two triskelia that meet to form a ring. In nsP3-ΔMD, two examples illustrate triskelion stability (top) and ring formation (bottom). In nsP3-ΔHVD, the ring formation occurs more slowly than in the other cases. **(e)** Two-dimensional network formation of nsP3-wt, nsP3-ΔMD and nsP3-ΔHVD. Representative frames from t= 0s to t*∼*300s from an HS-AFM video on the flat mica surface. **(f)** Diameter of rings in the three constructs studied imaged by HS-AFM on mica. Statistical significance was assessed by one-way ANOVA (*, p < 0.0001; **p = 0.0061). **(g)** Height distribution of the arms in the 2D network assembly of the constructs of nsP3. ***Statistical significance was assessed by one-way ANOVA with *p* < 0.0001.

At later stages of assembly, differences among the constructs became apparent compared to nsP3-wt. While the AUD domain alone did not assemble any organized structure (Fig. 5.b), under the removal of MD and the HVD domains the assembly took place (Fig. 5.c and Fig. 5d). Yet, they showed altered shape and dynamics at the different hierarchical levels: **(1)** *Triskelia*: Abundant in nsP3-wt and nsP3-ΔMD and presenting characteristic straight arms, these structures were scarce in nsP3-ΔHVD and their arms were more curved (Fig. 5.c, circles) as shown by the quantitative analysis of a 190-frame HS-AFM sequence (Suppl. Movie 5). The triskelia were significantly more abundant in nsP3-wt (21.9 ± 8.2 triskelia/frame) than in nsP3-ΔHVD (11.2 ± 3.5 triskelia/frame), while nsP3-ΔMD showed an intermediate abundance (13.8 ± 5.9 triskelia/frame (Suppl. Fig. 8.a and 8.b). The temporal evolution of triskelion abundance also showed the highest increase rate for nsP3-wt, followed by nsP3-ΔMD and nsP3-ΔHVD (Suppl. Fig. 8.c). **(2)** *Rings and other closed geometries (Fig. 5.d)*: The outer diameter differed significantly (Fig. 5.f and Suppl. Fig. 9). HS-AFM images show that it increased 20% for nsP3-ΔMD (17.5 ± 4.1 nm, N=40, one-way ANOVA p < 0.0001) and by 10% for nsP3-ΔHVD (16.2 ± 3.1 nm, N=33, one-way ANOVA, p = 0.0061) compared to nsP3-wt (14.7 ± 2.1 nm, N=54). The inner diameter was conserved in all cases (nsP3-wt: 8.7 ± 1.4 nm, N = 54; nsP3-ΔMD: 9.4 ± 2.3 nm, N = 40; nsP3-ΔHVD: 8.8 ± 1.6 nm; N = 33, Fig. 5.f), which suggests that the AUD defines the inner ring. The role of the AUD in defining the core of the HS is consistent with cryo-EM data that shows that the AUD forms structural core of these tubes ^5^. **(3)** *2D network*: all the constructs assembled 2D networks on the mica surface (Fig. 5.e). Yet, the absence of the MD produces an increase of the diameter of the arms of the 2D network of 19% with respect the nsP3-wt, which does not occur in the absence of the HVD (Fig. 5.g, 2D network height: nsP3-wt, 4.1 ± 0.5 nm (N=21); nsP3-ΔMD, 5.2 ± 0.4 nm (N=13); nsP3-ΔHVD, 4.1 ± 0.5 nm (N=21), one-way ANOVA, p < 0.01). Hence, the MD seems to act as a belt which tightens the arms of the nsP3 2D network.

In conclusion, the AUD constitutes the fundamental unit required for proper HS assembly, and the MD and HVD domains modulate nsP3 assembly preventing nonspecific aggregation, improving solubility and promoting the formation of well-defined intermediates. The HVD creates a furry envelope, but is also essential for the formation of triskelion with straight arms. While the MD is a mechanical belt that compacts the HS.

### Domain deletions. Structural and Mechanical Regulation of the Helical Scaffolds by MD, HVD, and AUD

To complement our data on the early stages and gain further insight on the roles of the MD and HVD domains, we expanded our examination by domain deletions to the late stages of the assembly the helical scaffolds. HSs were assembled in the absence of the MD or HVD domain and studied by NS-TEM, cryo-EM, and HS-AFM (following the same approach as we did in the early stages). It was determined that:

**(1)** *HVD Controls the Level of Aggregation, but MD does not.* Using NS-TEM, first, we examined the agregation of HS from precipitates formed just after affinity chromatography. We found the presence of HSs at elevated level of stacking and bundling (Fig. 6.a right top), in particular for nsP3-ΔHVD, which also showed the highest levels of precipitation. After purification of the remaining soluble nsP3-ΔHVD by size exclussion, we could not assemble any HSs possibly because of the low protein concentrations yielded (Fig. 6.a right bottom). In contrast, when we assessed the nsP3-ΔMD HS in the precipitates and in the purified solution, HSs were found in both cases with similar behaviour than the nsP3-wt (Fig. 6a left top and bottom, respectively). Hence, MD is not determinant in the aggregation and rather plays a structural role in scaffold organization whilst the HVD is a regulator of the HS bundling.
**(2)** *The MD acts as a Mechanical Belt Controlling Scaffold Geometry*. Representative HS-AFM images of the HSs of the nsP3-wt and nsP3-ΔMD are shown in Fig. 6.b. Quantitative analysis of HS-AFM images of purified nsP3-ΔMD HSs revealed a 41% increase in diameter with respect the wild type (from 34 ± 2 nm, N=30 to 48 ± 5 nm, N=18, mean ± SD, one-way ANOVA, p < 0.0001, Fig. 6.c), the finding indicates that the MD acts as a mechanical belt that constricts the scaffold. An additional cryo-EM structural analysis at 3.3 Å resolution (FSC 0.143) of the inner structure of nsP3-ΔMD HSs (Supp. Fig. 10, PDB: 9TJS, EMD-56018 ^22^, Table 1) confirmed that the MD does not alter the core helical parameters (helical rise of 2.89 Å, helical twist of -164.22°) or monomer-monomer interface area with respect the nsP3-wt HSs (1259 Å² in wt vs. 1298 Å² in ΔMD). Also, the final volumes were completely overlapping and no changes in interaction between monomers within helical structure were found, as the total interface area for one AUD monomer in nsP3-wt HSs was equal to 1259Å against 1298Å for nsP3-MD HSs, and the inner diameter (∼100 Å) across constructs preserved. Overall, the data supports that the MD’s primary function is to modulate the peripheral architecture, its radial expansion and flexibility of the scaffold (Fig. 6.e), rather than the core structure.
**(3)** *Mechanical Consequences of MD Deletion*. Unlike other tubular biological structures protofilaments ^23-24^, the HSs disassemble during HS-AFM imaging at minimal applied forces of tens of piconewtons ^21^. We profited from the progressive disassembly of the HSs to assess their mechanical stability. We measured the disassembly time of the HS for the different constructs of nsP3. Under the same mechanical perturbation (at imaging conditions of applied force ∼50 pN, 1.6μs/tap, 1 frame/second, 500 nm^2^ and 200x200 pixels), we found that the wild type HS takes 313% longer times to disintegrate that the HS of nsp3-ΔMD. Although cantilever spring constant may influence disassembly kinetics, the approximately fourfold difference was consistently reproduced across independent measurements (nsP3-wt, 62 ± 34 s, N=30; nsP3-ΔMD, 15 ± 8 s, N=18, mean ± SD, one-way ANOVA, p < 0.0001 Fig. 6.d). These results indicate that the MD substantially increases the mechanical stability of the HS under the imaging conditions used

**Figure 6.**
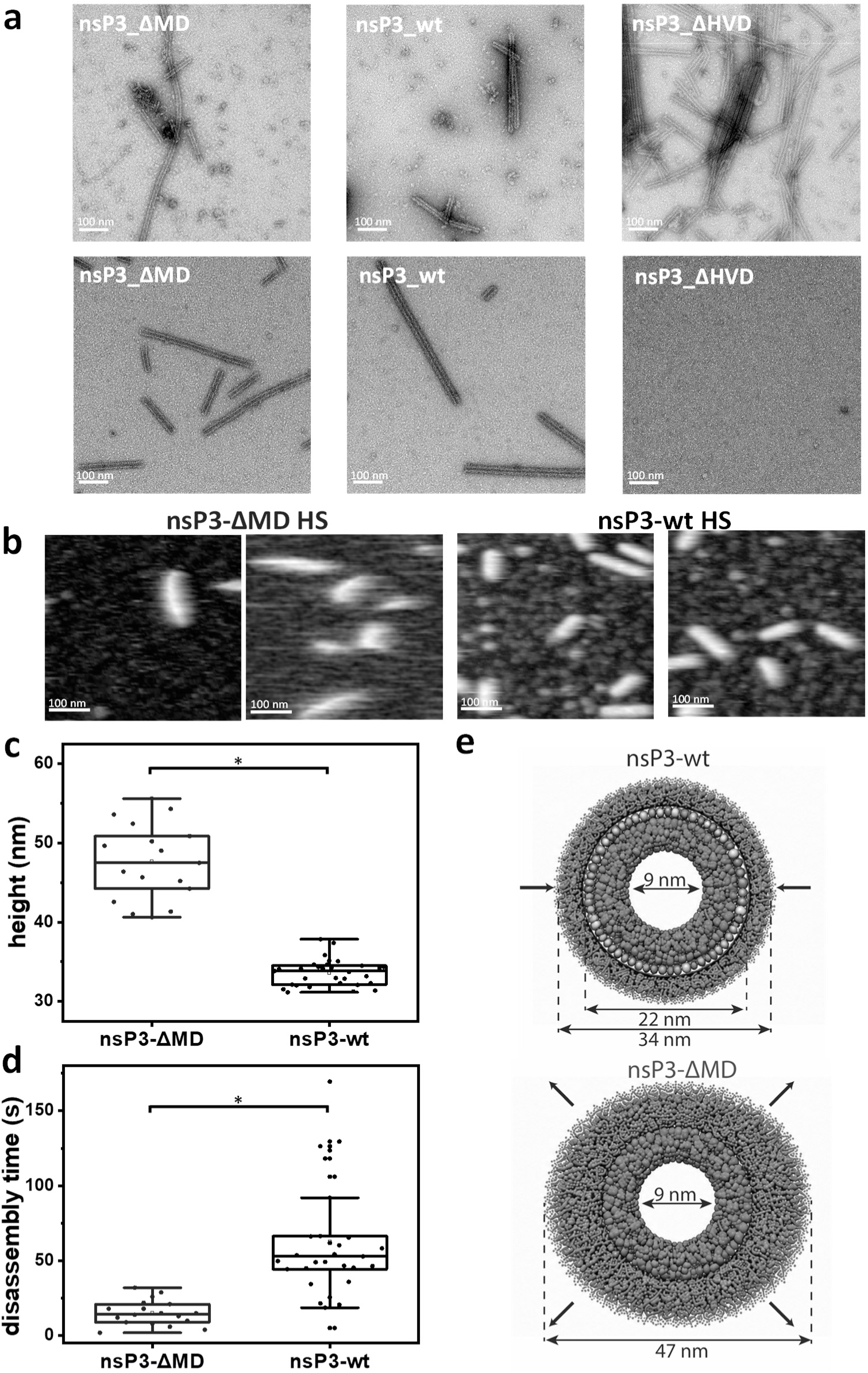
nsP3 in vitro helical scaffold formation and its deletion constructs. **(a) Top**, representative images of pellets formed after protein purification by affinity chromatography from cell lysates. **Bottom**, HSs fraction of purified with size exclusion chromatography proteins after overnight dialysis in low salt buffer (50 mM NaCl). **(b)** HS-AFM visualization of the HSs of nsP3-ΔMD (left) and nsP3-wt (right) adsorbed on mica. **(c)** Height distribution of nsP3-ΔMD (blue) and nsP3-wt (black) HSs. **(d)** Disassembly time distribution of nsP3-ΔMD (blue) and nsP3-wt (black) HSs. *Statistical significance in the two graphs was assessed by one-way ANOVA with *p* < 0.0001. **(e)** Role of the Macrodomain as a mechanical belt in nsP3 HSs. Schematic representation of the nsP3-wt and nsP3-ΔMD HSs illustrating the proposed effect of the MD on HVD deployment, it summarizes the structural model inferred from HS-AFM measurements: In nsP3-wt, the MD forms a constraining belt around the AUD core, limiting radial expansion of the flexible HVD and resulting in an overall HS diameter of 34 nm, with an inner diameter of 9 nm and an AUD–MD region extending to 22 nm. In contrast, deletion of the MD allows greater radial expansion and deployment of the HVD, resulting in an increased overall diameter of 47 nm while maintaining the 9 nm inner diameter.

The observed differences in the assembly of the HSs of nsP3-ΔHVD or nsP3-ΔMD determined that while the MD increases the cohesion of the HSs, the HVD prevents excessive levels of aggregation of the HSs.

## DISCUSSION

Recent work by Kril *et al.* (2024) ^5^ demonstrated that the Chikungunya virus nsP3 protein self-assembles into tubular helical scaffolds within the cytoplasm of infected and transfected cells. These HSs are both integral components of viral membrane associated replication complexes and of cytoplasmic Alphagranules by self organization into honeycomb-like arrays of HSs that can reach micrometric lengths. Cryo-EM analyses revealed that the HS core is formed by the helically organized alphavirus unique domain (AUD), while the macrodomain (MD) and hypervariable domain (HVD) project outward. However, the cryo-EM structures provided limited insight into the dynamic and mechanical roles of these external domains.

Here, by combining HS-AFM and EM, we provide complementary dynamic and mechanical information on nsP3 HSs, their intermediate assemblies, and the contribution of individual domains to their structure and stability. Our data reveal that the HSs form flexible, wool-like tubular architectures that lose the perfect helical symmetry seen in cryo-EM reconstructions. The measured diameters are larger than those previously reported, consistent with the presence of dynamic, flexible outer layers composed of the MD and HVD.

By mapping the height and topographic profiles of nsP3 HSs, we distinguish three concentric structural regions: (i) a central channel of ∼9 nm in diameter, (ii) a dense region of ∼22 nm corresponding to the AUD core and part of the MD, and (iii) an extended region of ∼34 nm encompassing the external, flexible HVD. The truncation of the MD (nsP3-ΔMD) increases the overall HS diameter to ∼47 nm, indicating that the MD acts as a mechanical belt that restricts the radial expansion and full deployment of the HVD (Fig. 6.e). The MD could retain the full deployment of HVD by holding the HVD in the interstitial space between MD and AUD along the HS (Fig. 3) and by direct interactions between MD and HVD as previously reported (see Krill et al. 2024 ^5^, Fig. 1.d) or additional interactions at the exterior of the MD belt. In the cellular context, the deployment of the HVD might be regulated by interactions with host or viral cofactors such as G3BP or nsP2 proteins, which accumulate with nsP3 HSs within alphagranules ^5^ or by BIN1 which accumulates in replication organelles ^25^. Post-translational modifications, as the reported phosphorylation of the HVD ^26-27^, can also regulate the deployment of the HVD, suggesting a sequential regulation of interactions through a controlled deployment of the HVD within alphagranules and replication organelles.

From a mechanical perspective, the HS-AFM recordings reveal that the MD domain confers rigidity and structural integrity to the HSs. Deletion of the MD notably reduces the resistance of HSs to tip-induced disassembly by shortening the disassembly time by approximately fourfold. Together, these findings establish that the MD stabilizes the HS both longitudinally and radially, acting as a mechanical stiffener. Beyond its known enzymatic activity, the MD thus serves a structural function essential for HS integrity in alphagranules and likely also in replication complexes.

HS-AFM movies also show the stepwise organization of nsP3 assemblies, providing a dynamic view of the early stages of HS formation. nsP3 monomers first associate into arcs and triskelion-like structures, which merge into ring-shaped intermediates and, subsequently, into polygons and 2D networks that precede tubular HS formation. Similar stepwise assembly pathways have been described for other proteins forming helical architectures, whose early intermediates have been directly visualized by HS-AFM, including tubulin protofilaments ^23^ and ESCRT-III tubules ^24^.

The high structural heterogeneity, dynamics and spatial segregation observed in the HS-AFM movies are consistent with the behaviour expected for intrinsically disordered protein-containing assemblies and biomolecular condensates, where transient and heterogeneous oligomeric states can coexist, creating multifunctional roles in viral processes ^28-29^. Our classification of the observed intermediates was therefore based on recurrent morphological features identified across the HS-AFM movies, rather than on the assumption of a strictly sequential assembly pathway. The assignment of monomers, oligomers, triskelia, and rings reflects characteristic and repeatedly observed morphologies, even though individual structures may persist for only a few frames.

Across all constructs, the internal diameter of the rings remains constant, indicating that the AUD defines the conserved inner core geometry. The absence of either the MD or the HVD does not prevent HS formation but alters the frequency, stability, and mechanical properties of intermediate assemblies. The HVD appears to prevent excessive aggregation of HSs, while the MD ensures the robustness of the tubes. These mechanistic insights build upon previous cellular and virological studies demonstrating the essential role of nsP3 helical scaffolds in alphagranule organization and viral replication, by revealing how these biologically relevant assemblies are formed and mechanically stabilized at the molecular level. nsP3 overexpression experiments in cells show how the expression of the AUD alone results in the generation of long filaments instead of defined pseudo-spherical granules. Even the deletion of MD or HVD result in a tendency to generate filaments or alter the granule size (see supplementary data Fig 9 A in Krill et al. 2024 ^5^ and PMID 22837213 ^30^). These phenotypes suggest that the physical and mechanical properties conferred by MD and HVD allows the proper formation of alphagranules in the cell, besides the important enzymatic and protein interaction roles, respectively, which are critical for infection.

Altogether, our results support a hierarchical model in which the AUD provides the structural core of the helical scaffold, the MD functions as a molecular belt that regulates rigidity and HVD deployment, and the HVD contributes flexibility, mediates external interactions with host factors and controls the level of bundling of the HSs. The balance between the stiffness imposed by the MD and the flexibility conferred by the HVD underlies the architecture and dynamics of the Alphagranule network. This integrated mechanical and structural framework highlights how nsP3 orchestrates both the formation and the stability of viral replication assemblies, revealing principles probably common to all togaviruses that may also extend to other homologous RNA virus replication complexes combining macrodomains and hypervariable regions (i.e. hepatitis E virus).

## MATERIAL AND METHODS

### EXPRESSION AND PURIFICATION OF CHIKV NSP3 AND ITS DELETIONAL CONSTRUCTS FROM INSECT CELLS

nsP3 and its deletional constructs were expressed in Hi5 cells (Thermo Fisher). The coding sequence of nsP3 (corresponding to residues 1334-1863 of the non-structural polyprotein sequence from the S27 African prototype, UniProt identifier: Q8JUX6) was synthesized by Gen9 as a codon-optimized gene and subcloned into a pFastBac vector (Thermo Fisher) with a C-terminal hexahistidine tag. For deletion of gene regions coding corresponding domains on nsP3 a Q5® Site-Directed Mutagenesis Kit was used with pre-selected primers. The deletions were verified by Sanger sequencing on Illumina platform (GeneWiz from Azenta Life Science). Recombinant bacmids and viruses were produced in YFP-DH10 Bac cells and Sf21 cells (Thermo Fisher), respectively, according to the manufacturer’s protocols. For protein expression, Hi5 cells in suspension at a density of 0.5–1.0 × 106 ml^−1^ were infected with baculoviruses at 1% volume of the culture. Cells were collected at 2–3 days after cell arrest.

Cells were resuspended in buffer A (20 mM Tris pH 8.0, 500 mM NaCl, 2 mM TCEP, 5% (v/v) glycerol, 1 mM PMSF and 20 mM imidazole) supplemented with, 10 μg × ml−1 DNase, 2 mM MgSO4, and complete, EDTA-free Protease Inhibitor Cocktail Tablets, and lysed by sonication. Lysates were centrifuged at 18,000g to remove unbroken cells and debris, and supernatants containing nsP3 and its deletion constructs were used for Ni-affinity purification on ӒKTA pure™ chromatography system with 5mL Cytiva HisTrap™ FF Columns. Resin was washed with 5 column volumes of buffer A and the proteins eluted over 1.5 column volumes with buffer A containing 500 mM imidazole. Elution fractions containing the protein were pooled and concentrated (Amicon 30k MW cut-off) for size-exclusion chromatography. Samples were applied to a Superdex 200 Increase 10/300 GL (GE Healthcare) pre-equilibrated with 20 mM Tris pH 8.0, 500 mM NaCl, 5% (v/v) glycerol and 2 mM TCEP. Fractions were analysed by SDS–PAGE to assess purity.

### NEGATIVE-STAINING TRANSMISION ELECTRON MICROSCOPY

The central fraction of the peaks corresponding to nsP3 or its deletion construct monomers were selected for analysis by NS-TEM. The proteins were diluted up to 0.2 mg × ml−1 and dialysed overnight at 4°C in buffer B (20 mM Tris pH 8.0, 50 mM NaCl, 5% (v/v) glycerol and 2 mM TCEP). For negative staining, 3.5 μl of protein was applied to Cu 300 mesh carbon grids (Agar Scientific), previously glow-discharged for 30 sec at 15 mA using a PELCO EasiGlow cleaning system (Ted Pella). Grids were blotted, washed 4 times with buffer B and stained with 1% (w/v) uranyl acetate before blotting and air drying. Images were acquired using a FEI Tecnai G2 Spirit microscope operating at 120 kV and equipped with a Veleta CCD camera (Olympus).

### GRID PREPARATION FOR CRYO-EM IMAGING

For nsP3-wt, quantifoil grid R2/1 was treated with plasma glow discharge (Quorum) two times for 60 s at 25 mA on each side. 2.0 μl of purified protein at 3 mg/ml in freezing buffer (20 mM Tris-HCl pH8.0, 50 mM NaCl, and 2 mM TCEP) were applied on the one side of the prepared grid, and 1.0 µl on another side. The grid was transferred in Leica GP plunge freezer and then blotted for 18.0 s from the back side. Grid was plunged into liquid ethane and then transferred and stored in liquid nitrogen until the imaging session. For data collection on nsP3-wt grids we used the Glacios 200kV microscope in Marseille, France. Vitrified grids were loaded into the microscope equipped with a Falcon 4i direct electron detector and a Selectris X energy filter. Cryo-EM data were acquired with EPU software (FEI) at a nominal magnification of ×160.000, with a pixel size of 0.71 Å and a total dose of 40 e/Å2 (see Table 1). 7168 micrographs were acquired with a flux of 8.8 electrons per square-pixel, fractionated into 40 frames with a defocus range from −0.6 μm to −3.0 μm. Whole data processing was conducted using the cryoSPARC.

For nsP3-ΔMD, quantifoil grid R2/2 was treated with plasma glow discharge (Quorum) two times for 60 s at 25 mA on each side, then the grid was covered with graphene oxide. 4.0 μl of purified protein at 0.8 mg/ml in freezing buffer (20 mM Tris-HCl pH8.0, 50 mM NaCl, and 2 mM TCEP) were applied on the prepared grid and left to incubate for 10 min for binding at RT. The grid was transferred in Vitrobot Mark IV plunge freezer and then blotted for 7.0 s. Grid was plunged into liquid ethane and then transferred and stored in liquid nitrogen until the imaging session. For data collection on nsP3-ΔMD grids we used the Glacios 200kV microscope at the PBSIM facility in Marseille, France. Vitrified grids were loaded into the microscope equipped with a Falcon 4i direct electron detector and a Selectris X energy filter. Cryo-EM data were acquired with EPU software (FEI) at a nominal magnification of ×130.000, with a pixel size of 0.90 Å and a total dose of 40 e/Å2 (see Table 1). 4610 micrographs were acquired with a flux of 14.1 electrons per square-pixel, fractionated into 40 frames with a defocus range from −0.6 μm to −3.0 μm. Whole data processing was conducted using the cryoSPARC.

### CRYO-EM IMAGE PROCESSING AND MODEL REFINEMENT

Acquired micrographs were imported into cryoSPARC for motion-correction and CTF estimation. After exposure curation 5916 micrographs for nsP3-wt and 4371 micrographs for nsP3-ΔMD remained. Combining two rounds of template free filament tracing and 2D classification on 100 micrographs, we obtained an initial set of templates for template-based filament tracing. Filament tracing was performed with a segment diameter of 300 Å/100 Å and a distance of 30 Å/28.9 Å based on measurements within 2D classes and resulted in 1.344.624/1.239.420 particle images for nsP3-wt/nsP3-ΔMD accordingly. 1.184.077/1.142.931 particles were extracted with an extraction crop size of 640/416 pixels and 2D-classified into 50 classes. Several consecutive rounds of 2D classification reduced the selection to 922.377/734.826 particles, only 338.875/325.402 particles were used for helical reconstruction.

Helical twist and rise estimates were used as in *Kril et al. 2024* ^5^. Helical parameters were refined during 3D reconstruction and converged to a twist of -164.170 degrees and a helical rise of 2.900 Å for nsP3-wt, and to a twist of -164.170 degrees and a helical rise of 2.891 Å for nsP3-ΔMD. The map refined to a nominal 2.77/3.31 Å resolution (according to 0.143 threshold) and a B-factor of 117.6/161.7. The predicted model of AUD from *Kril et al. 2024* ^5^ fitted perfectly in the maps. A model with five AUD including one protomer and four neighbour contacting protomers was done using Coot and real-space refined using Phenix on the 2.77/3.31 Å resolution map (PDB: 9TJR/9TJS and EMD: 56017/56018).

### SAMPLE PREPARATION FOR HIGH-SPEED ATOMIC FORCE MICROSCOPY

Freshly cleaved mica was used as the imaging substrate. A 2 μL aliquot of protein sample was deposited onto the mica surface in imaging buffer containing 20 mM Tris-HCl (pH 8.0), 2 mM TCEP, and either 500 mM NaCl or 50 mM NaCl, depending on the experiment. Samples were incubated on the mica for 20 min to allow protein adsorption. The surface was then gently washed with imaging buffer to remove non-adsorbed protein, after which 80 μL of the same buffer was added to the liquid cell for HS-AFM imaging. For experiments monitoring the assembly of nsP3 monomers, the following protein concentrations were used: nsP3-wt, 28 μg/mL; nsP3-ΔMD, 28 μg/mL; nsP3-AUD, 41 μg/mL; and nsP3-ΔHVD, 27 μg/mL. For imaging preassembled helical scaffolds (HSs), protein concentrations of 90 μg/mL (nsP3-wt) and 99 μg/mL (nsP3-ΔMD) were used.

### HIGH-SPEED ATOMIC FORCE MICROSCOPY

HS-AFM movies were acquired in amplitude modulation mode using high-resolution imaging parameters optimized for biomolecular systems, with a modified high-speed AFM (SS-NEX, RIBM, Tsukuba, Japan) ^14^. The system is equipped with a custom-built digital high-speed lock-in amplifier implemented on a reconfigurable FPGA (National Instruments, Austin, USA) and controlled via LabView. HS–AFM observations were performed in intermittent contact using short cantilevers designed for HS–AFM with a length of 6 μm, a nominal spring constant of 0.2 – 1.1 N/m, a resonance frequency of 600 kHz and a quality factor of 2 in solution that bear an electron beam deposition tip at the end of the cantilever (NanoWorld, Switzerland, ref. USC-F1.5-k0.6). The sensitivity of the AFM system to probe deflection was 0.1 V.nm^-1^. Because the cantilever oscillates at resonance in amplitude-modulation mode, the tip–sample interaction force varies throughout each oscillation cycle and cannot be defined by a single value ^31^. Therefore, rather than calibrating the imaging conditions based on the nominal spring constant of each cantilever, we standardized the oscillation amplitude relative to the thermal fluctuation baseline of the cantilever. The free oscillation amplitude was set to approximately 4 times the thermal fluctuation amplitude, providing sufficient signal-to-noise while minimizing the tip–sample interaction. During imaging, the oscillation amplitude was reduced by 10% relative to the free amplitude (90% setpoint). The free amplitude was around 10 Å. Under these conditions, the applied force by the HS-AFM tip on the sample was estimated to be tens of pN ^21^. The calibration of the height was performed using a diffraction gratting of parallel ridges of height 55 nm and pitch 278 nm (TDG01, NT-MDT, The Netherlands).

### HS-AFM IMAGE POST-TREATMENT AND ANALYSIS

The acquired HS-AFM images were plane-adjusted and flattened. The processing was limited to XY drift correction and a first-order adjustment of the X line. The raw images, were processed using the software tools WSxM ^32^, ImageJ ^33^, and Gwyddion ^34^.

### DLS MEASUREMENTS

nsP3-wt was diluted to 0.5 mg/ml in either buffer A (20 mM Tris-HCl (pH 8.0), 500 mM NaCl, 5% glycerol) or buffer B (20 mM Tris-HCl (pH 8.0), 50 mM NaCl, 5% glycerol) to a final volume of 60 µl. The samples were analysed using a DLS machine (Zetasizer Nano S) in automatic mode at 25°C during 30 measurement points. The experiment was performed in triplicate for each salt concentration, and the mean and standard deviation were calculated. For each time point, the hydrodynamic radius (R_h_) was determined as the mean size of the first peak in the DLS measurements.

### COMPARISON OF HS-AFM TOPOGRAPHY WITH STRUCTURAL MODELS

To compare the HS-AFM topography to the cryo-EM structural data, a custom-made code was was implemented using the macro coding language of free available software and plugins (Fiji is just ImageJ, the version 1.54f 64-bit works correctly). The procedure is the following: 1) the HS-AFM movie was opened in Fiji as .tif file. 2) the frame of interest is selected. 3) the code of the Macro01_AFM2Chimera_ (available in Suppl. Information) is run. 3) the code Macro02_AFM2Chimera_ (available in Suppl. Information) is run. 5) The file created by the Macro02_AFM2Chimera is opened in USCF Chimera (version 1.13.1 works correctly), the map values are inverted, a volume filter of 2 applied, and the range adjusted until a full surface obtained. Finally, the scale bar and the voxel coordinates are used to ajust the dimensions of the molecules in the USCF Chimera surface visualization to that of the HS-AFM topography. 6) The resulting surface map is saved in .mrc format.

In ChimeraX, the nsP3 ring with 12 subunits was obtained by applying a 12 fold symmetry to the nsP3 monomer (PDB: 8PK7). The model was fitted to the AFM volumes in ChimeraX manually, next fine-adjusted using ChimeraX Fit in Map utility. Additionally, the PMID 40133775, AFlecto, a software for the modelling of intrinsic protein disordered regions was used.

The detailed procedure and source codes for the two macros, which can be run in Fiji/ImageJ, are provided in the Supplementary Information and https://github.com/DyNaMo-INSERM/AFM2Chimera_tiff2mrc.

## ACKNOWLEDGEMENTS

**N.M.-G.** was funded by the Margarita Salas Fellowship, from the Ministry of Universities (Spain), for the requalification of the Spanish University System, with funds from the European Union – NextGenerationEU (CA1/RSUE/2021-00868). Structural biology work has been carried out in the PBSIM platform for Cryo EM and biophysics. The PBSIM is supported by French Infrastructure for Integrated Structural Biology (FRISBI) ANR-10-INBS-0005. At the AFMB. **A.K.** is supported by a CENTURI PhD fellowship financed by "Investissements d’Avenir" program Excellence Initiative of Aix-Marseille Université - A*MIDEX (ANR-16-CONV-0001), as well as the Fondation pour la Recherche Médicale 4^th^ year of thesis program (FDT202504020438). **J.R.** ’s team was supported by the Agence Nationale de la Recherche (ANR-22-CE11-0023-01). **I.C.**’s team was supported by the Agence Nationale de la Recherche (ANR-21-CE42-0031).

## AUTHOR CONTRIBUTIONS

J.R. and I.C. designed the study. N.M-G. performed HS-AFM. A.K. produced the proteins and perfomed the EM experiments. N.M-G and I.C. analysed the HS-AFM data. A.K and J.R. analysed the EM data. All the authors wrote the paper.

## COMPETING INTERESTS

The authors declare no competing interests.

## DATA, CODE, AND MATERIALS AVAILABILITY

All data and code needed to evaluate and reproduce the results in the paper are present in the paper and/or the Supplementary Materials.

This study did generated new materials.

The raw HS-AFM data obtained and analyzed in this study, as well as the data used in the graphs presented in the figures, are publicly available in the Zenodo repository at [DOI: 10.5281/zenodo.22231364].

The Fiji-Macro code to compare the HS-AFM topography to the cryo-EM structural data is available at https://github.com/DyNaMo-INSERM/AFM2Chimera_tiff2mrc.

The electron density maps of nsP3-wt and nsP3-dMD obtained and analyzed in this study are publicly available in Electron Microscopy Data Bank (EMDB) with codes EMD-56017 and EMD-56018 respectively. The pentamers of AUD fitted within these maps are publicly available in RCSB Protein Data Bank (RCSB PDB) with codes 9TJR and 9TJS.

## Notes

https://www.rcsb.org/structure/unreleased/9TJS

https://github.com/DyNaMo-INSERM/Particles-maximal-height-movie-histogram

### Competing Interest Statement

The authors have declared no competing interest.

## REFERENCES

1. Kril, V.; Aïqui-Reboul-Paviet, O.; Briant, L.; Amara, A., New Insights into Chikungunya Virus Infection and Pathogenesis. Annual review of virology 2021, 8 (1), 327–347.

2. Chikungunya vaccine approved. Nature biotechnology 2023, 41 (12), 1667.

3. Jones, R.; Bragagnolo, G.; Arranz, R.; Reguera, J., Capping pores of alphavirus nsP1 gate membranous viral replication factories. Nature 2021, 589 (7843), 615–619.

4. Zhang, K.; Law, Y. S.; Law, M. C. Y.; Tan, Y. B.; Wirawan, M.; Luo, D., Structural insights into viral RNA capping and plasma membrane targeting by Chikungunya virus nonstructural protein 1. Cell host & microbe 2021, 29 (5), 757–764.e3.

5. Kril, V.; Hons, M.; Amadori, C.; Zimberger, C.; Couture, L.; Bouery, Y.; Burlaud-Gaillard, J.; Karpov, A.; Ptchelkine, D.; Thienel, A. L.; Kümmerer, B. M.; Desfosses, A.; Jones, R.; Roingeard, P.; Meertens, L.; Amara, A.; Reguera, J., Alphavirus nsP3 organizes into tubular scaffolds essential for infection and the cytoplasmic granule architecture. Nat Commun 2024, 15 (1), 8106.

6. Tan, Y. B.; Chmielewski, D.; Law, M. C. Y.; Zhang, K.; He, Y.; Chen, M.; Jin, J.; Luo, D., Molecular architecture of the Chikungunya virus replication complex. Sci Adv 2022, 8 (48), eadd2536.

7. Laurent, T.; Kumar, P.; Liese, S.; Zare, F.; Jonasson, M.; Carlson, A.; Carlson, L. A., Architecture of the chikungunya virus replication organelle. eLife 2022, 11.

8. Meshram, C. D.; Agback, P.; Shiliaev, N.; Urakova, N.; Mobley, J. A.; Agback, T.; Frolova, E. I.; Frolov, I., Multiple Host Factors Interact with the Hypervariable Domain of Chikungunya Virus nsP3 and Determine Viral Replication in Cell-Specific Mode. Journal of virology 2018, 92 (16).

9. Foy, N. J.; Akhrymuk, M.; Akhrymuk, I.; Atasheva, S.; Bopda-Waffo, A.; Frolov, I.; Frolova, E. I., Hypervariable domains of nsP3 proteins of New World and Old World alphaviruses mediate formation of distinct, virus-specific protein complexes. Journal of virology 2013, 87 (4), 1997–2010.

10. Götte, B.; Liu, L.; McInerney, G. M., The Enigmatic Alphavirus Non-Structural Protein 3 (nsP3) Revealing Its Secrets at Last. Viruses 2018, 10 (3).

11. Li, Z. Q.; Zhao, L. X.; Wang, S. Y.; Hu, C. Y.; Wang, Y. Y.; Yang, Y., YBX1 is required for assembly of viral replication complexes of chikungunya virus and replication of multiple alphaviruses. Journal of virology 2025, 99 (2), e0201524.

12. Shin, G.; Yost, S. A.; Miller, M. T.; Elrod, E. J.; Grakoui, A.; Marcotrigiano, J., Structural and functional insights into alphavirus polyprotein processing and pathogenesis. Proceedings of the National Academy of Sciences of the United States of America 2012, 109 (41), 16534–9.

13. Abraham, R.; Hauer, D.; McPherson, R. L.; Utt, A.; Kirby, I. T.; Cohen, M. S.; Merits, A.; Leung, A. K. L.; Griffin, D. E., ADP-ribosyl-binding and hydrolase activities of the alphavirus nsP3 macrodomain are critical for initiation of virus replication. Proceedings of the National Academy of Sciences of the United States of America 2018, 115 (44), E10457–e10466.

14. Ando, T.; Kodera, N.; Takai, E.; Maruyama, D.; Saito, K.; Toda, A., A high-speed atomic force microscope for studying biological macromolecules. Proceedings of the National Academy of Sciences 2001, 98 (22), 12468–12472.

15. Casuso, I.; Redondo-Morata, L.; Rico, F., Biological physics by high-speed atomic force microscopy. Philosophical Transactions of the Royal Society A 2020, 378 (2186), 20190604.

16. Villarrubia, J. S., Algorithms for Scanned Probe Microscope Image Simulation, Surface Reconstruction, and Tip Estimation. Journal of research of the National Institute of Standards and Technology 1997, 102 (4), 425–454.

17. Ando, T., Functional Implications of Dynamic Structures of Intrinsically Disordered Proteins Revealed by High-Speed AFM Imaging. Biomolecules 2022, 12 (12).

18. Ortega-Esteban, A.; Pérez-Berná, A. J.; Menéndez-Conejero, R.; Flint, S.; Martin, C. S.; De Pablo, P., Monitoring dynamics of human adenovirus disassembly induced by mechanical fatigue. Scientific reports 2013, 3 (1), 1434.

19. Rodríguez-Espinosa, M. J.; Rodríguez, J. M.; Castón, J. R.; de Pablo, P. J., Mechanical disassembly of human picobirnavirus like particles indicates that cargo retention is tuned by the RNA–coat protein interaction. Nanoscale Horizons 2023, 8 (12), 1665–1676.

20. Roos, W. H.; Radtke, K.; Kniesmeijer, E.; Geertsema, H.; Sodeik, B.; Wuite, G. J., Scaffold expulsion and genome packaging trigger stabilization of herpes simplex virus capsids. Proceedings of the National Academy of Sciences 2009, 106 (24), 9673–9678.

21. Umeda, K.; Kodera, N., Excitation frequency dependence of noise and minimum detectable force in amplitude-modulation atomic force microscopy. Physical Review Applied 2026, 25 (1), 014033.

22. EMD-56018, https://www.ebi.ac.uk/emdb/EMD-56018. 2025.

23. Keya, J. J.; Inoue, D.; Suzuki, Y.; Kozai, T.; Ishikuro, D.; Kodera, N.; Uchihashi, T.; Kabir, A. M. R.; Endo, M.; Sada, K.; Kakugo, A., High-Resolution Imaging of a Single Gliding Protofilament of Tubulins by HS-AFM. Sci Rep 2017, 7 (1), 6166.

24. Chiaruttini, N.; Redondo-Morata, L.; Colom, A.; Humbert, F.; Lenz, M.; Scheuring, S.; Roux, A., Relaxation of Loaded ESCRT-III Spiral Springs Drives Membrane Deformation. Cell 2015, 163 (4), 866–79.

25. Neuvonen, M.; Kazlauskas, A.; Martikainen, M.; Hinkkanen, A.; Ahola, T.; Saksela, K., SH3 domain-mediated recruitment of host cell amphiphysins by alphavirus nsP3 promotes viral RNA replication. PLoS pathogens 2011, 7 (11), e1002383.

26. Thaa, B.; Biasiotto, R.; Eng, K.; Neuvonen, M.; Götte, B.; Rheinemann, L.; Mutso, M.; Utt, A.; Varghese, F.; Balistreri, G.; Merits, A.; Ahola, T.; McInerney, G. M., Differential Phosphatidylinositol-3-Kinase-Akt-mTOR Activation by Semliki Forest and Chikungunya Viruses Is Dependent on nsP3 and Connected to Replication Complex Internalization. Journal of virology 2015, 89 (22), 11420–37.

27. Teppor, M.; Žusinaite, E.; Merits, A., Phosphorylation Sites in the Hypervariable Domain in Chikungunya Virus nsP3 Are Crucial for Viral Replication. Journal of virology 2021, 95 (9).

28. Rawlinson, S. M.; Chakraborty, S.; Sethi, A.; David, C. T.; Harrison, A. R.; Bird, L. E.; Rozario, A. M.; Uthishtran, S.; Ardipradja, K.; Zhao, T., Conformational dynamics, RNA binding, and phase separation regulate the multifunctionality of rabies virus P protein. Nature Communications 2025, 16 (1), 9491.

29. Brocca, S.; Grandori, R.; Longhi, S.; Uversky, V., Liquid–liquid phase separation by intrinsically disordered protein regions of viruses: Roles in viral life cycle and control of virus–host interactions. International journal of molecular sciences 2020, 21 (23), 9045.

30. Fros, J. J.; Domeradzka, N. E.; Baggen, J.; Geertsema, C.; Flipse, J.; Vlak, J. M.; Pijlman, G. P., Chikungunya virus nsP3 blocks stress granule assembly by recruitment of G3BP into cytoplasmic foci. Journal of virology 2012, 86 (19), 10873–10879.

31. Stan, G.; King, S. W., Atomic force microscopy for nanoscale mechanical property characterization. Journal of Vacuum Science & Technology B 2020, 38 (6).

32. Colom, A.; Derivery, E.; Soleimanpour, S.; Tomba, C.; Molin, M. D.; Sakai, N.; González-Gaitán, M.; Matile, S.; Roux, A., A fluorescent membrane tension probe. Nature chemistry 2018, 10 (11), 1118–1125.

33. Schindelin, J.; Arganda-Carreras, I.; Frise, E.; Kaynig, V.; Longair, M.; Pietzsch, T.; Preibisch, S.; Rueden, C.; Saalfeld, S.; Schmid, B., Fiji: an open-source platform for biological-image analysis. Nature methods 2012, 9 (7), 676–682.

34. Nečas, D.; Klapetek, P., Gwyddion: an open-source software for SPM data analysis. Open Physics 2012, 10 (1), 181–188.

